# Flow environment and matrix structure interact to determine spatial competition in *Pseudomonas aeruginosa* biofilms

**DOI:** 10.1101/077354

**Authors:** Carey D. Nadell, Deirdre Ricaurte, Jing Yan, Knut Drescher, Bonnie L. Bassler

## Abstract

Bacteria often live in biofilms, which are microbial communities surrounded by a secreted extracellular matrix. Here, we demonstrate that hydrodynamic flow and matrix organization interact to shape competitive dynamics in *Pseudomonas aeruginosa* biofilms. Irrespective of initial frequency, in competition with matrix mutants, wild type cells always increase in relative abundance in straight-tunnel microfluidic devices under simple flow regimes. By contrast, in microenvironments with complex, irregular flow profiles - which are common in natural environments - wild type matrix-producing and isogenic non-producing strains can coexist. This result stems from local obstruction of flow by wild-type matrix producers, which generates regions of near-zero flow speed that allow matrix mutants to locally accumulate. Our findings connect the evolutionary stability of matrix production with the hydrodynamics and spatial structure of the surrounding environment, providing a potential explanation for the variation in biofilm matrix secretion observed among bacteria in natural environments.

**Impact Statement:** The feedback between hydrodynamic flow conditions and biofilm spatial architecture drives competition in *P. aeruginosa* biofilms, and can explain the variation in biofilm production observed among bacteria in natural environments.

## Introduction

In nature, bacteria predominantly exist in biofilms, which are surface-attached or free-floating communities of cells held together by a secreted matrix [1–3]. The extracellular matrix defines biofilm structure, promoting cell-cell and cell-surface adhesion and conferring resistance to chemical and physical insults [4–9]. The matrix also plays a role in the population dynamics of biofilm-dwelling bacteria [2, 10–15]; simulations and experiments using *Vibrio cholerae, Pseudomonas fluorescens*, and *Pseudomonas aeruginosa* show that matrix-secreting cell lineages can smother or laterally displace other cell lineages and, in so doing, outcompete neighboring matrix non-producing cells [16–24]. However, not all wild and clinical isolates of these species produce biofilm matrices, despite the clear ecological and competitive benefits of possessing a matrix [25–27]. Fitness trade-offs between the benefits of being adhered to surfaces and the ability to disperse to new locations can cause variability in matrix production [17, 26], but is not known whether selective forces within the biofilm environment itself might drive the coexistence of strains that make matrix with strains that do not. Here we explore how selection for matrix production occurs in biofilms on different surface geometries and flow regimes, including those that are relevant inside host organisms and in abiotic environments such as soil.

The local hydrodynamics associated with natural environments can have dramatic effects on biofilm matrix organization. This phenomenon has been particularly well established for *P. aeruginosa*, a common soil bacterium [28] and opportunistic pathogen that thrives in open wounds [29, 30], on sub-epithelial medical devices [31], and in the lungs of cystic fibrosis patients [32–36]. Under laminar flow in simple microfluidic channels, *P. aeruginosa* forms biofilms with intermittent mushroom-shaped tower structures [32, 37–39]. Under irregular flow regimes in more complex environments, however, *P. aeruginosa* also produces sieve-like biofilm streamers that protrude into the liquid phase above the substratum [40–43]. These streamers – whose structure depends on the secreted matrix – are proficient at catching cells, nutrients, and debris that pass by, leading to clogging and termination of local flow [44]. The spatial and temporal characteristics of flow thus combine to alter matrix morphology, which, in turn, feeds back to alter local hydrodynamics and nutrient advection and diffusion [11, 45, 46]. By modifying community structure and solute transport in and around biofilms [47], this feedback could have a significant influence on the evolutionary dynamics of matrix secretion in natural environments.

Here, we examined the relationship between flow regime and matrix-mediated biofilm competition using strains of *P. aeruginosa* PA14 that differ only in their production of Pel, a visco-elastic matrix polysaccharide that is central to biofilm and streamer formation [37, 44, 48, 49]. Using a combination of fluid flow visualization and population dynamics analyses, we reveal a novel interaction between hydrodynamic conditions, biofilm architecture, and competition within bacterial communities.

## Results/Discussion

We performed competition experiments with wild type PA14 and an otherwise isogenic strain deleted for *pelA*, which is required for synthesis of Pel [50]. Deletion of *pelA* significantly impairs biofilm formation in PA14, which does not naturally produce Psl, an additional matrix polysaccharide secreted by other *P. aeruginosa* isolates [51, 52]. Wild type cells produced GFP, and Δ*pelA* mutants produced mCherry. Experiments in shaken liquid culture using genetically identical wild type cells producing GFP or mCherry confirmed that fluorescent protein expression constructs had no measurable effect on growth rate (Figure S1). Our first goal was to compare the population dynamics of the wild type and Δ*pelA* strains in typical straight-chamber microfluidic devices, which have simple parabolic flow regimes, and in porous environments containing turns and corners, which have irregular flow profiles and better reflect the packed soil environments that *P. aeruginosa* often occupies [28, 53, 54]. To approximate the latter environment, we used microfluidic chambers containing column obstacles; the size and spacing distributions of the column obstacles were designed specifically to simulate soil or sand. Analogous methods have been used previously to study bacterial growth [55] and the behavior of *Caenorhabditis elegans* [56] in realistic environments while maintaining accessibility to microscopy. In our setup, flow was maintained through these chambers at rates comparable to those experienced by *P. aeruginosa* in a soil environment [57].

### The flow regime alters selection for matrix production in biofilms

Several approaches are available to study how competitive dynamics differ in particular flow environments. Most commonly, one would monitor biofilm co-cultures of wild type and Δ*pelA* PA14 cells over time until their strain compositions reached steady state. Performing such time-series experiments was not possible here due to a combination of low fluorescence output in the early phases of biofilm growth coupled with phototoxicity incurred by cells during epifluorescence imaging. To circumvent this issue, we measured the change in frequency of wild type and Δ*pelA* cells as a function of their initial ratio in both straight-tunnel chambers and soil-mimicking chambers containing column obstacles. From these measurements, we could infer the final stable states of Pel-producing and non-producing cells as a function of surface topography and flow conditions. This method is commonly used to measure the behavior of dynamical systems, and it has been employed successfully in a variety of analogous experimental applications [17, 18, 23, 58–60].

In straight-tunnel chambers with simple parabolic flows, wild type PA14 increased in relative abundance regardless of initial population composition, indicating uniform positive selection for Pel secretion. This result is consistent with recent studies of *V. cholerae* and *Pseudomonas* spp. demonstrating that the secreted matrix cannot be readily exploited by non-producing mutants (Figure 1A,B) [17–23]. Confocal microscopy revealed that Pel-producers mostly excluded non-Pel-producing cells from biofilm clusters in straight tunnel chambers (Figure 2A,B), although some Δ*pelA* mutants resided on the periphery of wild type biofilms. The liquid effluent from these chambers contained an over-representation of the Δ*pelA* mutant relative to wild type, consistent with the interpretation that Δ*pelA* strain was displaced from the substratum over time (Figure 2C). When wild type and Δ*pelA* cells competed in microfluidic devices simulating porous microenvironments, by contrast, there was a pronounced shift to negative frequency-dependent selection for Pel production (Figure 1A,C). Wild type PA14 was selectively favored at initial frequencies below ~0.6. Above this critical frequency, the Δ*pelA* mutant was favored. From this result, we can infer that in this porous environment, Δ*pelA* null mutants can grow and stably coexist with wild type Pel-producers.

**Figure 1.**
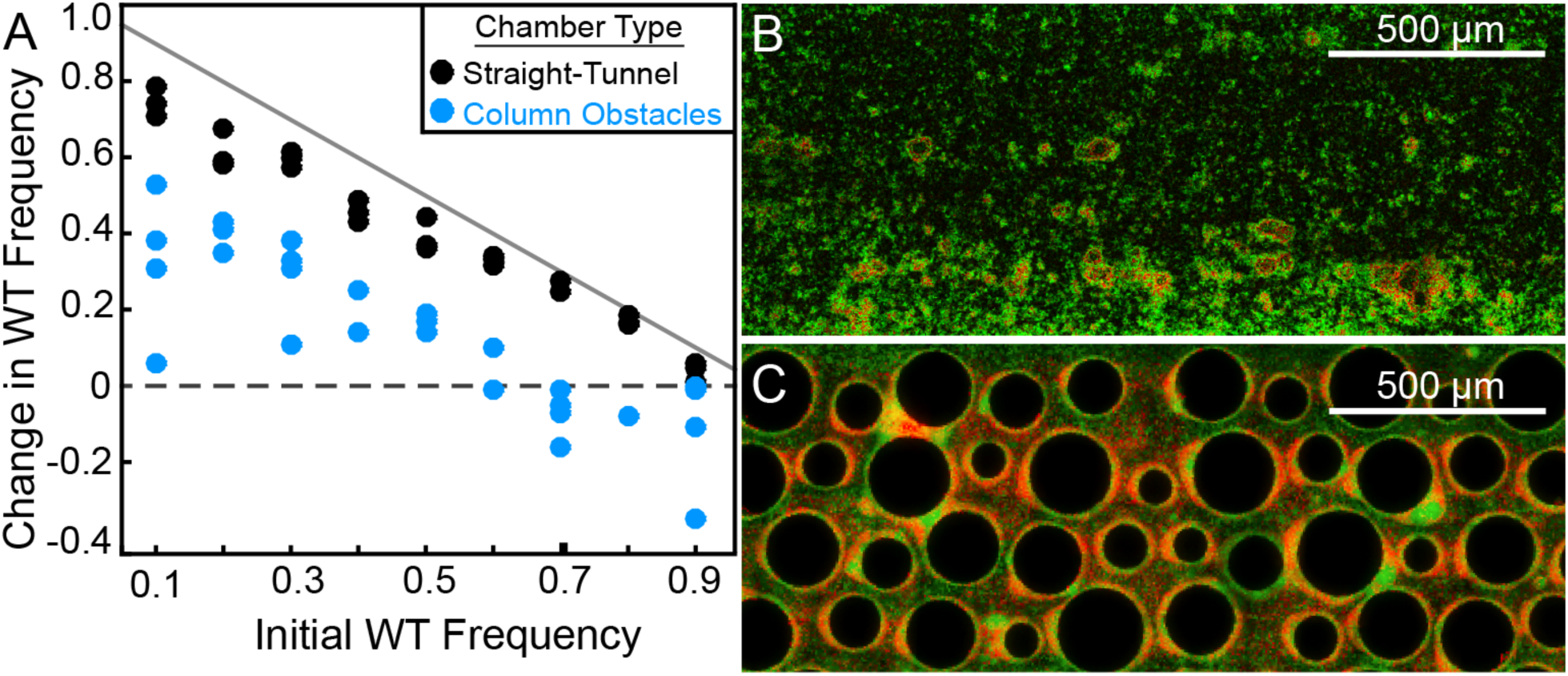
Wild type *P. aeruginosa* PA14 outcompetes the Δ*pelA* mutant under simple flow conditions, but the two strains coexist under complex flow conditions. (A) Wild type and Δ*pelA* strains were co-cultured at a range of initial frequencies in the simple straight-tunnel (black data) or column-containing (blue data) microfluidic chambers. In both cases, fresh minimal M9 medium with 0.5% glucose was introduced at flow rates adjusted to the cross-sectional area of each chamber type to equalize the average initial flow velocity in both environments. The diagonal gray line denotes the maximum possible increase in wild type frequency for a given initial condition. Each data point is an independent biological replicate. (B) A maximum intensity projection (top-down view) of a confocal *z*-stack of wild type (green) and Δ*pelA* (red) biofilms in simple flow chambers. (C) An epifluorescence micrograph (top-down view) of wild type (green) and Δ*pelA* (red) biofilms after 72 h growth in a flow chamber containing column obstacles to simulate a porous environment with irregular flows. Images in (B) and (C) were taken from chambers in which the wild type was inoculated at a frequency of 0.7.

**Figure 2.**
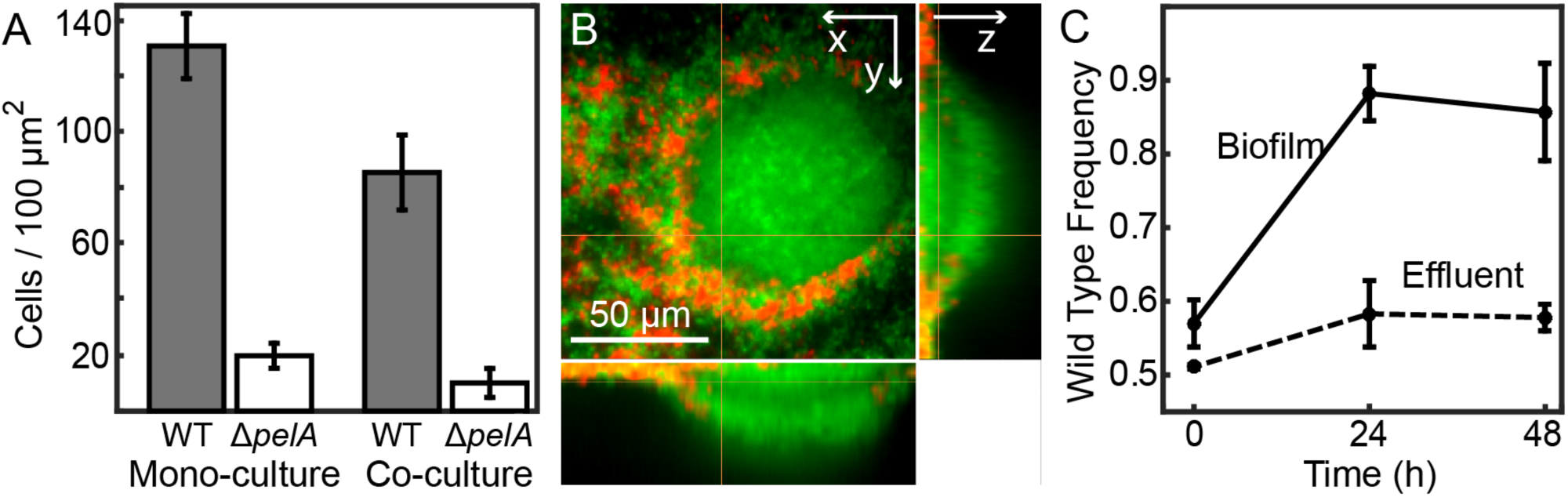
Matrix production confers a competitive advantage to wild type *P. aeruginosa* PA14 in biofilms under simple flow conditions. (A) Absolute abundances of wild type and Δ*pelA* strains in monoculture and co-culture in simple flow microfluidic chambers (Bars denote means +/− S.D. for *n* = 3-6). The two strains were inoculated alone (left two bars) or together at a 1:1 ratio (right two bars). (B) Single optical plane 3 μm from the surface, and *yz*- and *xz*-projections at right and bottom respectively, of the wild-type and Δ*pelA* strains growing in co-culture 48 h after inoculation at a 1:1 ratio. (C) Relative abundance of wild type and Δ*pelA* strains in the liquid effluent of straight-chamber microfluidic devices (Points denote means +/− S.D. for *n* = 3). Wild type and Δ*pelA* cells were combined at a 1:1 initial ratio and co-inoculated on the glass substratum of simple flow chambers. At 0, 24, and 48 h, 5 µL of effluent was collected from chamber outlets. Wild type frequency was calculated within biofilms and within the liquid effluent for each time point.

### Wild type P. aeruginosa obstructs porous environments, generating shear-free regions that Pel-deficient mutants can occupy

Previous work has shown that in environments with corners, wild type *P. aeruginosa* produces biofilm streamers that catch cells and other debris in the surrounding flow [44]. Such streamers were produced by the wild type in our system. Although Δ*pelA* cells could be found in these structures, they were not abundant, so cell capture by streamers cannot account for the observed coexistence of the two strains (Figure S2). This result suggests that streamers do indeed catch debris, consistent with prior studies [41, 44], but that in our system, Δ*pelA* cells are not present at high enough density in the passing liquid phase to accumulate substantial population sizes by this manner.

Our microscopy-based observation of chambers containing column obstacles suggested that wild type biofilms gradually obstructed some of the regions located between columns over time. We hypothesized that partial clogging could render those portions of the chambers more suitable for growth of the Δ*pelA* strain, which was previously shown to be sensitive to removal by shear [51, 52]. This hypothesis predicts that Δ*pelA* cells should be found predominantly in regions of the chamber that have been clogged by wild type biofilms, or downstream of such regions. To test this prediction, we repeated our co-culture competition experiment with wild type and Δ*pelA* cells in chambers containing columns, measured the distribution of each strain as above, and then introduced fluorescent beads to the chambers by connecting new influent syringes to the inflow tubing. By tracking the beads with high frame-rate microscopy, we could trace the presence and absence of fluid flow and then superimpose this information onto the spatial distributions of wild type and Δ*pelA* mutant cells (Figure S3). Wild type biofilms accumulated intermittently, often with clusters of Δ*pelA* cells in close proximity. Importantly, in support of our prediction, Δ*pelA* cells grew most successfully in regions that were locally obstructed by wild type biofilms (Figure 3A,B). Consistent with prior work [44], Δ*pelA* cells did not clog chambers when grown in isolation, supporting the interpretation that Δ*pelA* accumulation relies on and only occurs after clogging by the wild type. This result is consistent with prior indications that Pel-producers do indeed have a higher tolerance for shear than Pel-deficient cells [51, 52]. We directly confirmed this prediction in our system by independently growing each strain in isolation under varying shear stress in straight-tunnel chambers (Figure 3C).

**Figure 3.**
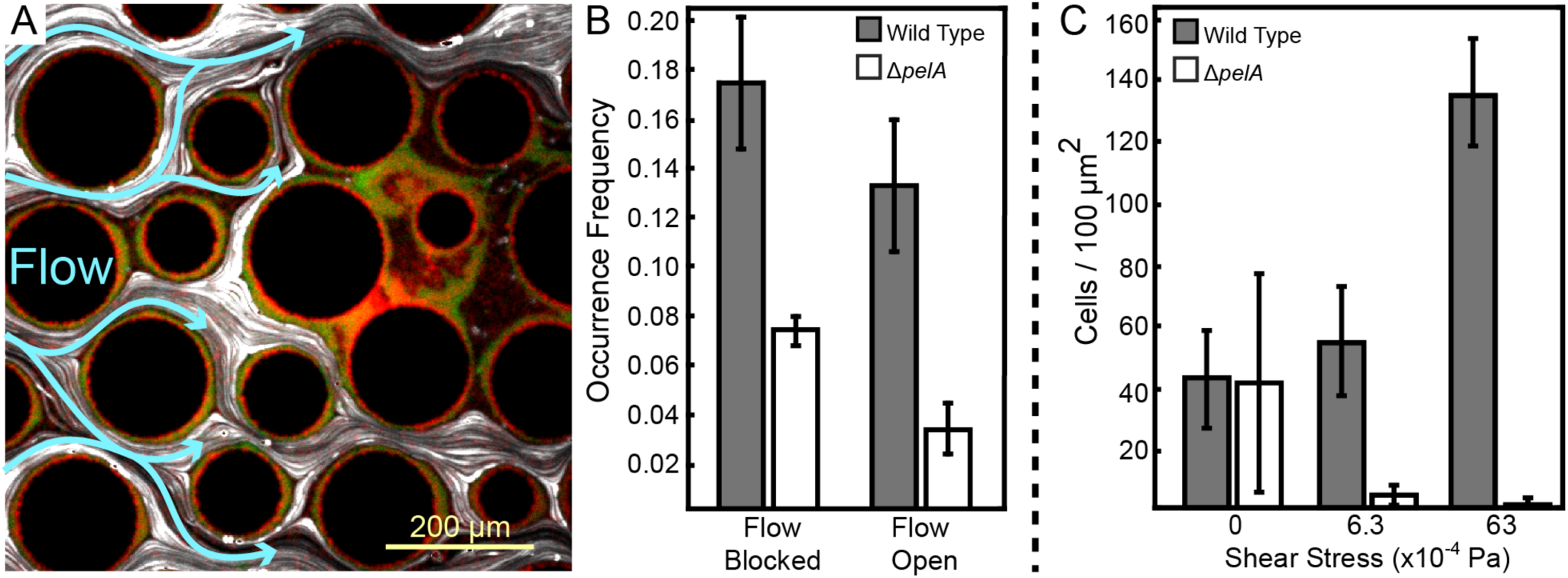
Pel-deficient mutants occupy locations protected from flow due to local clogging by wild type *P. aeruginosa* biofilms. (A) Wild type (green) and Δ*pelA* (red) *P. aeruginosa* strain mixtures were inoculated into complex flow chambers with irregularly-spaced column obstacles. Biofilms were imaged using confocal microscopy, after which fluorescent beads were flowed through the chamber. The presence or absence of flow was monitored through averaging successive exposures of bead tracks (white lines are bead tracks; blue arrows highlight flow trajectories). (B) Analysis of co-occurrence of flow and wild type or Δ*pelA* cell growth at the end of 1:1 competition experiments in complex flow chambers with column obstacles, as illustrated by the micrograph in (A). The occurrence of wild type (gray) and Δ*pelA* (white) cell clusters are shown as a function of whether local flow has been blocked or remains open after 72 h of competition (Bars denote means +/− S.E. for *n* = 3). There is no significant difference in wild type occurrence in the presence and absence of flow (two-sample t=0.995, *df*=4, p=0.376), but the Δ*pelA* strain is more significantly more likely to occur when flow is blocked at a p<0.05 threshold with Bonferroni correction for two pairwise comparisons (two-sample t=3.60, *df*=4, p=0.0227). (C) Biofilm growth of wild type *P. aeruginosa* PA14 (gray) and the Δ*pelA* mutant (white) in monoculture in flat, straight-tunnel flow chambers under different shear stress exposure treatments (Bars denote means +/− S.D. for *n* = 5-10).

Collectively, our findings suggest that when *P. aeruginosa* wild type and Δ*pelA* mutants experience irregular flow in heterogeneous environments, wild type biofilm formation causes partial clogging, regionally reducing local flow speed. The lack of flow generates favorable conditions for the Δ*pelA* strain, whose biofilm would otherwise be removed by shear forces, enabling it to proliferate locally using nutrients diffusing from the bulk liquid phase [61].

## Conclusions

Our results show that a feedback process operates between hydrodynamic flow conditions, biofilm spatial architecture, and competition in *P. aeruginosa* biofilms. Pel non-producers do not appear to directly exploit the matrix material produced by wild type cells, but they can take advantage of the low-shear conditions that wild type matrix producers generate in complex flow environments. *P. aeruginosa* is notorious as an opportunistic pathogen of plants and animals, including humans [62]. It also thrives outside of hosts, for example, in porous niches such as soil [63]. Despite the well-documented ecological benefits of matrix secretion during biofilm formation, environmental and clinical isolates of *P. aeruginosa* exhibit considerable variation in their production of matrix components, including loss or overexpression of Pel [25, 48]. Our results offer an explanation for natural variation in the ability of *P. aeruginosa* to produce extracellular matrix, particularly among bacteria in porous environments: the evolutionary stable states of extracellular matrix secretion vary with the topographical complexity of the flow environment in which bacteria reside.

## Materials and Methods

### Strains

All strains are derivatives of *Pseudomonas aeruginosa* PA14. Wild type PA14 strains constitutively producing fluorescent proteins [44] were provided by Albert Siryaporn (UC Irvine), and they harbor genes encoding either EGFP or mCherry under the control of the P_A1/04/03_ promoter in single copy on the chromosome [64]. The Δ*pelA* strain was constructed using the lambda red system modified for *P. aeruginosa* [65].

### Liquid growth rate experiments

To determine maximum growth rates and the potential for fluorescent protein production to cause fitness differences, bacterial strains were grown overnight in M9 minimal medium with 0.5% glucose at 37º C. Overnight cultures were back-diluted into minimal M9 medium with 0.5% glucose at room temperature and monitored until their optical densities at 600 nm were ~0.2, corresponding to logarithmic phase. Cultures were back diluted again into minimal M9 medium with 0.5% glucose and transferred to 96 well plates at room temperature. This experiment was repeated for 4 biological replicates (different overnight inoculation cultures), each repeated for 6 technical replicates (different wells within a 96 well plate). Measurements of culture optical density at 600 nm were taken once per 10 minutes until saturation, corresponding to stationary phase. MatLab curve fitting software was used to calculate the maximum growth rate of each strain. These experiments confirmed that the fluorescent protein markers had no measurable effect on growth rates and thus did not contribute to competitive outcomes in our experiments.

### Microfluidics and competition experiments

Microfluidic devices consisting of poly(dimethylsiloxane) (PDMS) bonded to 36 mm × 60 mm glass slides were constructed using standard soft photolithography techniques [66]. We used flat, straight-tunnel microfluidic devices with no obstacles to simulate environments with simple parabolic flow profiles, and we used devices with PDMS pillars interspersed throughout the chamber volume to simulate environments with complex (i.e. irregular, non-parabolic) flow profiles. The size and spatial distributions of these column obstacles were determined by taking a cross-section through a simulated volume of packed beads mimicking a simple soil environment. Flow rates through these two chamber types were adjusted to equalize the average initial flow velocities, although the local flow velocity within each chamber varied as biofilms grew during experiments (see main text).

For all competition experiments, bacterial strains were grown overnight. The following morning, aliquots of the overnight cultures were added to Eppendorf tubes, and their optical densities were equalized prior to preparation of defined mixtures of wild type and Δ*pelA* cells. 100 µL volumes of the wild type strain alone, the Δ*pelA* strain alone, or mixtures of the two strains (for competition experiments), were introduced into microfluidic chambers using 1 mL syringes and Cole-Parmer polytetrafluoroethylene tubing (inner diameter = 0.30 mm; outer diameter = 0.76 mm). After 3 h, fresh tubing connected to syringes containing fresh minimal M9 medium with 0.5% glucose were inserted into the inlet channels. The syringes (3 mL BD Syringe, 27G) were mounted onto high precision syringe pumps (Harvard Apparatus), which were used to tune flow rate according to empirical measurements of flow speeds in soil [57]. Biofilms were grown at room temperature. It should be noted that microfluidic experiments in the obstacle-containing chambers experience a high failure rate, in which no biofilms appear to grow after the 72 h period of the experiment. No data could be extracted from these chambers, which were omitted from analysis. This problem was overcome by performing the experiment at high replication in parallel. Sufficient data were thus collected to populate the relevant panels in Figures 1 and 3 of the main text. In the case of competition experiments, one replicate was defined as the output from one independently inoculated microfluidic chamber (e.g., Figure 1A). For experiments in which biofilm growth was measured as a function of flow-mediated shear stress (Figure 3C), one replicate was defined as the output from one imaging location within a microfluidic chamber, with 2-3 locations per chamber being sampled.

### Microscopy and image analysis

Mature biofilms were imaged using a Nikon Ti-E inverted microscope via a widefield epifluorescence light path (using a 10x objective) or a Borealis-modified Yokogawa CSU-X1 spinning disk confocal scanner (using a 60x TIRF objective). A 488 nm laser line was used to excite EGFP, and a 594 nm laser line was used to excite mCherry. Quantification of biofilm composition was performed using MatLab and Nikon NIS Elements analysis software [60]. Imaging of biofilms could only be performed once for each experiment, precluding time-series analyses, due to phototoxicity effects after multiple rounds of imaging. Phototoxicity was a particularly notable issue here due to dimness of the fluorescent proteins in *P. aeruginosa*, which made long exposures necessary to capture images of sufficient quality for later analysis. For this reason we opted for inferential population dynamics analysis as described in the main text.

### Effluent Measurements

To measure strain frequencies in the biofilm effluent of straight-tunnel chambers (Figure S2), 1:1 strain mixtures of wild type and Δ*pelA* cells were prepared and inoculated into simple flow chambers according to the procedure outlined above for competition experiments. At 0, 24, and 48 h, 5 µL samples were collected from the microfluidic chamber outlet tubing, mixed vigorously by vortex, and plated onto agar in serial dilution. After overnight growth at 37º C, plates were imaged with an Image Quant LAS 4000. Cy3 and Cy5 fluorescence settings were used for EGFP and mCherry excitation, respectively. Image Quant TL Colony Counting software was used to measure the relative abundance of each strain.

### Flow Tracking Experiments

1:1 mixtures of the wild type and the Δ*pelA* mutant were prepared and introduced into obstacle-containing flow chambers according to the procedure described above. Minimal M9 medium with 0.5% glucose was introduced into the chambers for 72 h as described above. The entire chamber was then imaged using widefield epifluorescence microscopy to document the locations of wild type and Δ*pelA* cell clusters. Subsequently, the influent syringes were replaced with syringes containing yellow-green fluorescent beads (Invitrogen; sulfate-modified, diameter = 2 µm) at a concentration of 0.3%, and beads suspensions were flowed into the microfluidic chambers. To determine the presence or absence of flow with respect to the spatial distributions of wild type and Δ*pelA* cells, and to obtain large images for statistics, the entire chamber was imaged with a 1 s exposure time, over which traveling beads were captured as streaks. It should be noted that this experiment also has a high failure rate do to the sensitivity of the microfluidic chambers to removal and re-insertion of syringes, and required optimization to execute successfully. Custom MatLab code was written to correlate the presence or absence of fluid flow with the accumulation of wild type and Δ*pelA* cells. In brief, the positions of the columns were first identified and used to divide the chamber into triangular sampling areas using a network structure in which columns served as nodes and straight lines between column centers served as edges. Within each sampling triangle, the area covered by columns was first removed, and subsequently, the averaged Δ*pelA* and wild type fluorescence intensities in the remaining area were used to determine if a region had wild type and/or Δ*pelA* accumulation. In parallel, each sampling area was scored for the presence of flow in the corresponding bead tracking images (Figure S3).

### Data Display and Statistical Tests

In all cases where displayed, bars denote the mean values of the measurements taken, and with the exception of Figures 3B and S1, the error bars denote standard deviations. For the data in Figure 3B, error bars denote standard errors, and we report the results of a two-tailed t-test comparing the wild type occurrence frequency in regions of soil-mimicking chambers where flow was obstructed, versus the wild type occurrence frequency in regions where flow was unobstructed. A second t-test was performed to make the same comparison for Δ*pelA* cells. The p-values from these tests were evaluated against a critical threshold of p<0.05 adjusted by Bonferroni correction for two pairwise comparisons. Two t-tests were also performed on data in Figure S1 measuring the maximum growth rates of our strains in liquid culture.

## Acknowledgments

We thank members of the B.L.B. laboratory, as well as Thomas Bartlett and Alvaro Banderas for helpful discussions. This work was supported by the Alexander von Humboldt Foundation (CDN), the Human Frontier Science Program Grant CDA00084/2015-C (KD), the Max Planck Society (KD), the Howard Hughes Medical Institute (BLB), NIH Grant 2R37GM065859 (BLB), National Science Foundation Grant MCB-0948112 (BLB), and National Science Foundation Grant MCB-1344191 (BLB).

## Supplemental Information

### Supplemental Materials and Methods

**Figure S1.**
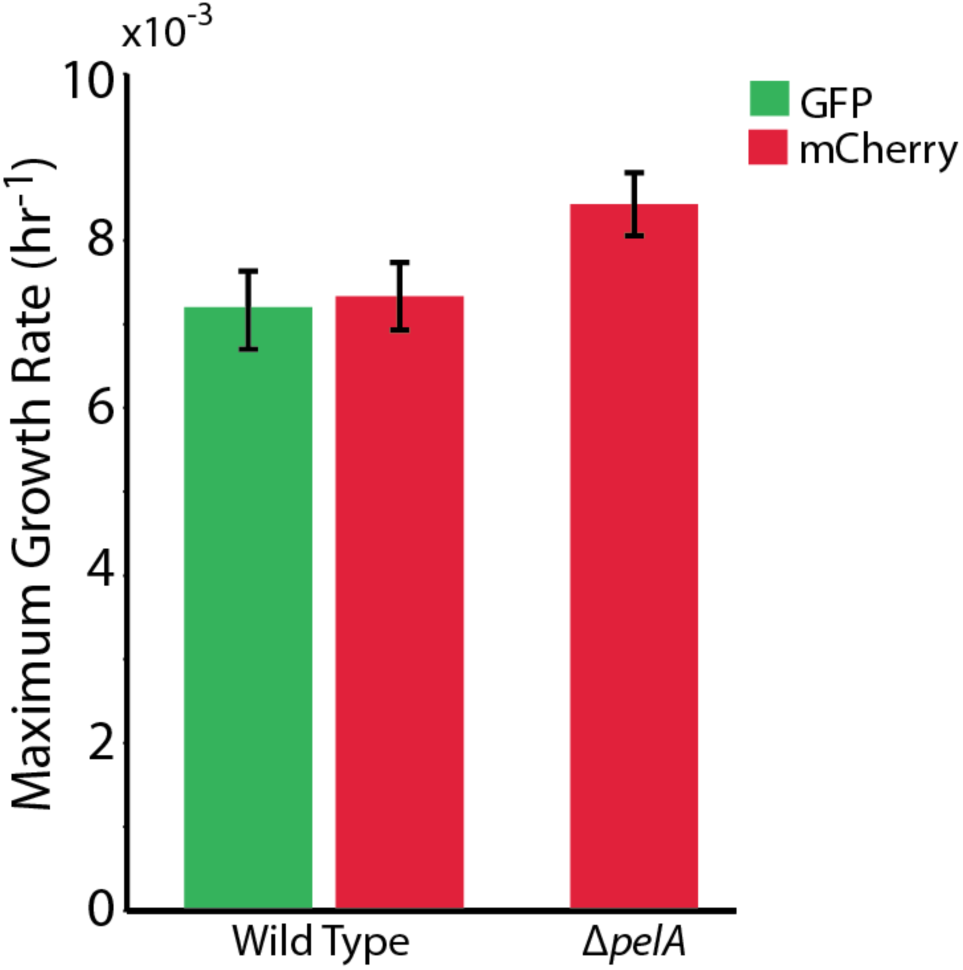
The maximum growth rates of *P. aeruginosa* PA14 wild type and Δ*pelA* cells in mixed liquid culture (M9 + 0.5% glucose). *n* = 4 biological × 6 technical replicates (different overnight cultures and microtiter plate wells, respectively (Bars denote means +/− S.E.). Growth rates were measured as the maximum slope of growth curves of cultures whose optical density at 600nm was taken every 30 minutes until stationary phase. Cultures were grown in minimal M9 medium with 0.5% glucose at ambient room temperature. The maximum growth rates of wild type cells expressing GFP and wild type cells expressing mCherry were not different, indicating that the fluorescent protein constructs did not introduce growth rate bias (two-sample t=0.235, *df*=6, p=0.822). A two-sample t-test comparing the maximum growth rate of the Δ*pelA* strain against the pooled data for wild type (t=2.31, *df*=10, p=0.0432) suggests a growth rate decrement on the part of wild type relative to Δ*pelA*, but was not statistically significant at a critical p<0.05 threshold with Bonferroni correction for two pairwise comparisons.

**Figure S2.**
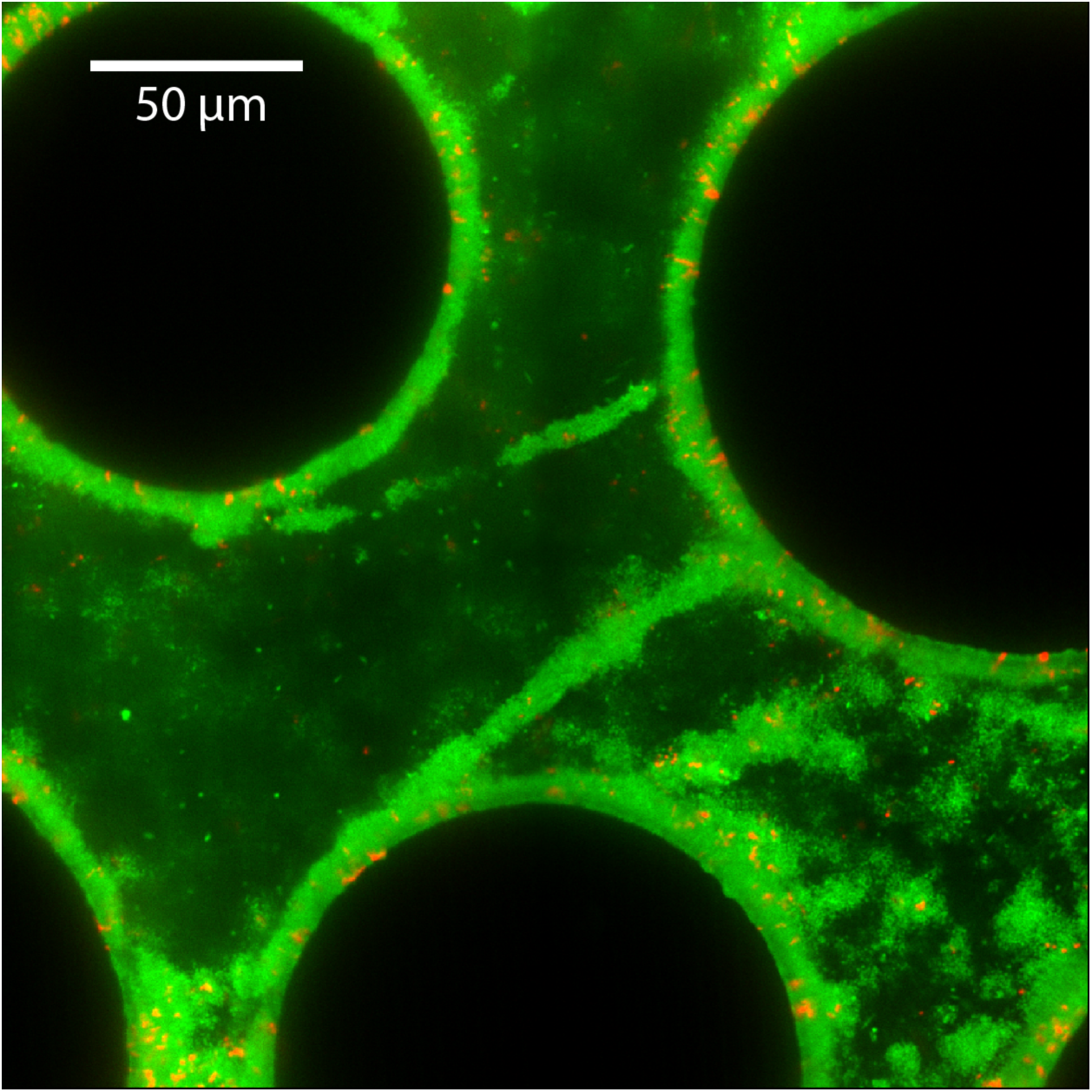
Streamer structures produced by wild type *P. aeruginosa* PA14 (green) in microfluidic chambers with complex flow profiles do not capture large numbers of co-cultured Δ*pelA* mutants (red) over 72 h of biofilm growth (black circles are column obstacles).

**Figure S3.**
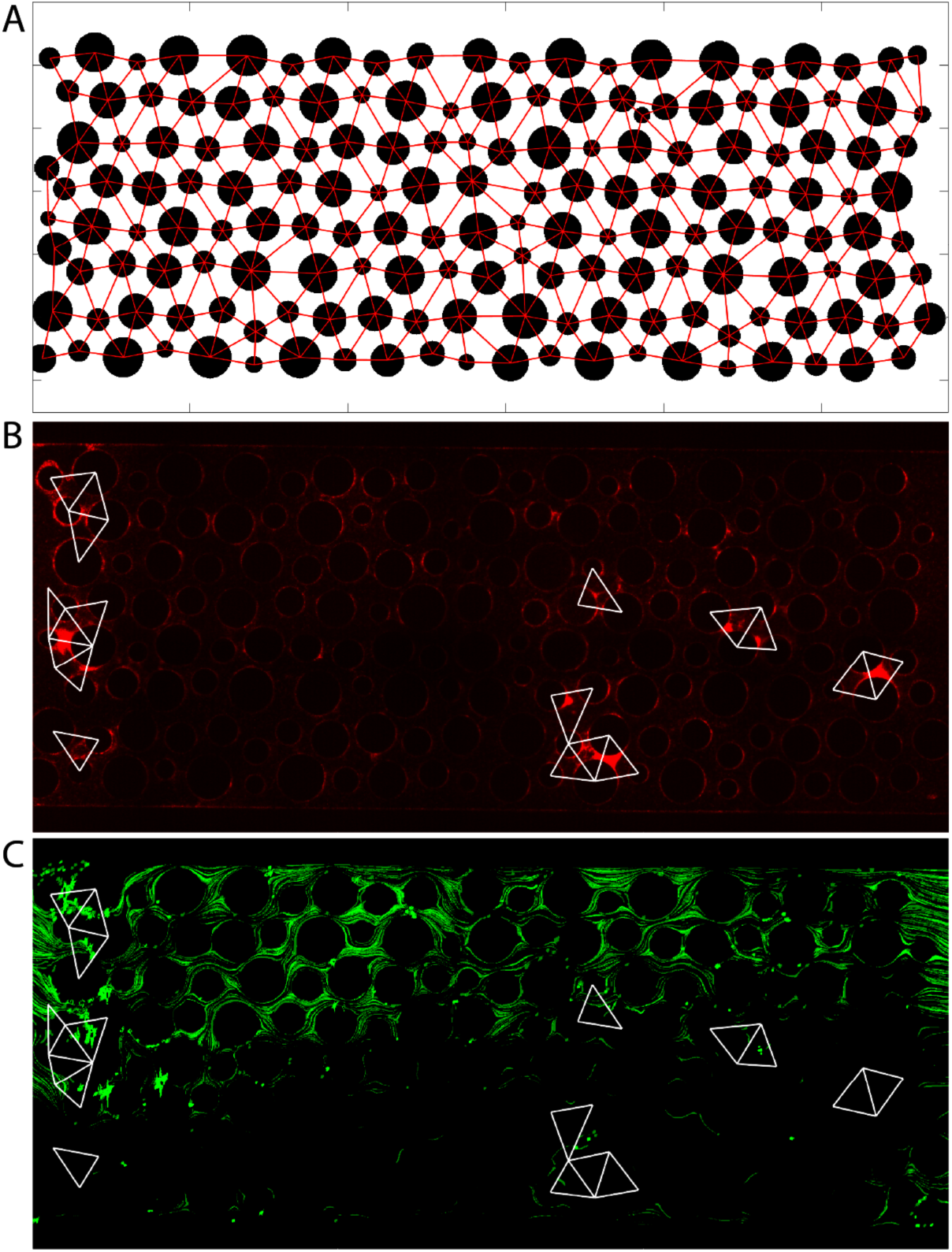
Analysis procedure for correlating local flow and accumulation of wild type and Δ*pelA* cells (only Δ*pelA* cells are shown in this figure for simplicity). (A) First, the positions of the columns were identified from the fluorescence image. The centers of the columns served as nodes to divide the spaces into triangular regions. Within each region, the area covered by the column was removed. (B) Averaged intensity within each sampling region was calculated, and a threshold was set to determine if biofilm accumulation occurred in each region. White triangles correspond to regions identified as containing biofilms using this method. (C) The corresponding image with fluorescent beads (green) to track local flows. Typically, 10 to 15 images were taken and integrated to cover the flow regions sampled by the beads. Overlaid are triangular regions showing Δ*pelA* cell accumulation (from (B)). Δ*pelA* cells accumulated primarily in regions lacking flow due to upstream clogging (bottom) or in regions immediately downstream of obstructed areas, indicated by accumulation of non-moving beads (left, (C)).

